# Tissue morphogenesis mediated by the Arabidopsis receptor kinase STRUBBELIG involves a clathrin-dependent process

**DOI:** 10.1101/448084

**Authors:** Jin Gao, Ajeet Chaudhary, Prasad Vaddepalli, Marie-Kristin Nagel, Erika Isono, Kay Schneitz

## Abstract

**Highlight:** The Arabidopsis receptor kinase STRUBBELIG is internalized by clathrin-mediated endocytosis and affects clathrin-dependent processes in a tissue-dependent manner.

**Abstract:** Signaling mediated by cell surface receptor kinases is central to the coordination of growth patterns during organogenesis. Receptor kinase signaling is in part controlled through endocytosis and subcellular distribution of the respective receptor kinase. For the majority of plant cell surface receptors the underlying trafficking mechanisms are not characterized. In Arabidopsis, tissue morphogenesis relies on the atypical receptor kinase STRUBBELIG (SUB). Here, we approach the endocytic mechanism of SUB. Our data reveal that a functional SUB:EGFP fusion is ubiquitinated in vivo. We further show that plasma membrane-bound SUB:EGFP becomes internalized in a clathrin-dependent fashion. We also find that SUB:EGFP associates with the *trans*-Golgi network and accumulates in multivesicular bodies and the vacuole. Coimmunoprecipitation experiments reveal that SUB:EGFP and clathrin are present within the same protein complex. Our genetic analysis shows that *SUB* and *CLATHRIN HEAVY CHAIN 2* promote root hair patterning. By contrast, *SUB* behaves as a negative regulator of a clathrin-dependent process during floral development. Taken together, the data indicate that SUB undergoes clathrin-mediated endocytosis, that this process does not dependent on stimulation of SUB signaling by an exogenous agent, and that *SUB* genetically interacts with clathrin-dependent pathways in a tissue-specific manner.

## Introduction

Intercellular communication is a central requirement for tissue morphogenesis as cells have to orchestrate their relative behavior to allow proper organ development. Cell surface receptor kinases play crucial roles in this process. Control of tissue morphogenesis in *Arabidopsis thaliana* involves the leucine-rich repeat (LRR) receptor kinase (RK) STRUBBELIG (SUB). SUB, also known as SCRAMBLED (SCM), controls several developmental processes, including floral morphogenesis, integument outgrowth, leaf development and root hair patterning (Chevalier *et al.*, 2005; Kwak *et al.*, 2005; Lin *et al.*, 2012). SUB represents an atypical receptor kinase as enzymatic activity of its kinase domain is not required for its function in vivo (Chevalier *et al.*, 2005; Vaddepalli *et al.*, 2011). SUB is glycosylated in the endoplasmic reticulum (ER) (Hüttner *et al.*, 2014), subject to ER-associated protein degradation (Vaddepalli *et al.*, 2011; Hüttner *et al.*, 2014), and found at the plasma membrane (PM) (Yadav *et al.*, 2008; Vaddepalli *et al.*, 2014). SUB not only localizes to the PM but is also present at plasmodesmata (PD), channels interconnecting most plant cells (Otero *et al.*, 2016; Sager and Lee, 2018), where it physically interacts with the PD-specific protein QUIRKY (QKY) (Vaddepalli *et al.*, 2014). In line with a function in RK-mediated control of PD-based intercellular communication *SUB* and *QKY* function in a non-cell-autonomous manner (Yadav *et al.*, 2008; Vaddepalli *et al.*, 2014) indicating that SUB signaling involves a yet unknown factor that moves between cells. More recently, a genetic link of SUB signaling to cell wall biology has also been put forward as the cell wall-localized β-1,3 glucanase ZERZAUST (ZET) participates in SUB signal transduction and *sub*, *qky* and *zet* mutants share overlapping defects in cell wall biochemistry (Fulton *et al.*, 2009; Vaddepalli *et al.*, 2017).

The maintenance of the PM composition is in part achieved through exocytosis/secretion and endocytosis (Paez Valencia *et al.*, 2016; Reynolds *et al.*, 2018). In general plant cells internalize PM-bound material or cargo via membrane transport into the trans-Golgi network (TGN), an organelle that also functions as an early endosome (EE) and that serves as a sorting hub for subsequent trafficking pathways. The cargo may get recycled back to the PM via secretory vesicles. Cargo may also be destined to degradation via endosomal transport to multivesicular bodies (MVBs), also known as late endosomes (LEs), containing intra-luminal vesicles (ILVs). MVBs eventually fuse with the tonoplast discharging their content into the vacuolar lumen where it becomes degraded.

Internalization of PM proteins is mediated by clathrin-dependent and clathrin-independent endocytosis (Geldner and Robatzek, 2008; Robinson *et al.*, 2008; Irani and Russinova, 2009; Paez Valencia *et al.*, 2016; Reynolds *et al.*, 2018). Clathrin-mediated endocytosis (CME) is a central mechanism for the internalization of PM-localized material or cargo (Dhonukshe *et al.*, 2007; Paez Valencia *et al.*, 2016; Reynolds *et al.*, 2018). CME involves the budding of cargo-containing clathrin-coated vesicles (CCVs) from the PM. CCVs consist of vesicles surrounded by a polyhedral lattice of clathrin triskelia being made of three clathrin heavy chains (CHCs), each bound by a clathrin light chain (CLC) (Fotin *et al.*, 2004). In Arabidopsis, three genes encode CLC chains while the likely redundant acting *CHC1* and *CHC2* encode clathrin heavy chains (Scheele and Holstein, 2002). Clathrin is also present at the TGN/EE, at a subpopulation of MVB/LEs, and at the cell plate indicating that it functions in multiple vesicular trafficking steps, and cytokinesis in the plant cell (Samuels *et al.*, 1995; Staehelin and Moore, 1995; Konopka *et al.*, 2008; Fujimoto *et al.*, 2010; Stierhof and El Kasmi, 2010; Kang *et al.*, 2011; Van Damme *et al.*, 2011; Ito *et al.*, 2012).

Fine-tuning the spatio-temporal dynamics of receptor-mediated endocytosis and endosomal trafficking is a central element in the regulation of RK-dependent signal transduction. Such a mechanism can for example maintain the steady-state level of RKs at the PM through recycling internalized RKs back to the PM, promote signaling by activated RK complexes localized on endosomes, or attenuate RK signaling by controlled removal of activated receptors from the PM followed by sorting into MVBs and finally degradation in the vacuole (Geldner and Robatzek, 2008; Irani and Russinova, 2009; Di Rubbo and Russinova, 2012; Bakker *et al.*, 2017; Critchley *et al.*, 2018).

Following their internalization and subsequent trafficking upon RK stimulation by exogenous application of ligand has been instrumental in analyzing the endocytic pathways of several plant RKs, including BRASSINOSTEROID INSENSITIVE 1 (BRI1) (Russinova *et al.*, 2004; Geldner *et al.*, 2007; Irani *et al.*, 2012; Di Rubbo *et al.*, 2013), FLAGELLIN SENSING 2 (FLS2) (Robatzek *et al.*, 2006; Beck *et al.*, 2012; Du *et al.*, 2013; Mbengue *et al.*, 2016), or PEP1 RECEPTOR 1 (PEPR1) (Ortiz-Morea *et al.*, 2016). No ligand for SUB has been described to date rendering such an experimental strategy currently impossible. However, some RKs undergo endocytosis independently of exogenous application of ligand, including BRI1 (Russinova *et al.*, 2004; Geldner *et al.*, 2007; Jaillais *et al.*, 2008), SOMATIC EMBRYOGENESIS RECEPTOR-LIKE KINASE 1 (SERK1) (Shah *et al.*, 2001; Shah *et al.*, 2002), BRI1-ASSOCIATED RECEPTOR KINASE 1 (BAK1)/SERK3 (Russinova *et al.*, 2004), and ARABIDOPSIS CRINKLY4 (ACR4) (Gifford *et al.*, 2005). SUB can be found in internal compartments as well (Kwak and Schiefelbein, 2008; Yadav *et al.*, 2008; Vaddepalli *et al.*, 2011; Wang *et al.*, 2016b) and it was recently shown that ovules of plants homozygous for a hypomorphic allele of *HAPLESS13* (*HAP13*) preferentially accumulate signal from a functional SUB:EGFP reporter in the cytoplasm, rather than the PM (Wang *et al.*, 2016b). *HAP13*/*AP1M2* encodes the µ1 subunit of the adaptor protein (AP) complex AP1 that is present at the TGN/EE network and is involved in post-Golgi vesicular trafficking to the PM, vacuole and cell-division plane (Park *et al.*, 2013; Teh *et al.*, 2013; Wang *et al.*, 2013). Interestingly, ovules of plants with reduced *HAP13*/*AP1M2* activity show *sub*-like integuments (Wang *et al.*, 2016b). These results indicate that the AP1 complex is involved in subcellular distribution of SUB in a functionally relevant manner.

Here, we have further assessed the internalization and subsequent endocytic trafficking behavior of SUB. We show that the intracellular domain of a functional SUB:EGFP fusion protein becomes ubiquitinated. Upon endocytosis SUB:EGFP is sorted to MVBs and the vacuole. Our data further indicate that CME contributes to internalization of SUB:EGFP. Finally, we provide genetic data suggesting that *CHC* is part of the *SUB*-dependent signaling mechanism that mediates root hair patterning while *SUB* is a negative regulator of a *CHC*-dependent pathway in floral tissue.

## Materials and methods

### Plant work, plant genetics and plant transformation

*Arabidopsis thaliana* (L.) Heynh. var. Columbia (Col-0) and var. Landsberg (*erecta* mutant) (L*er*) were used as wild-type strains. Plants were grown as described earlier (Fulton *et al.*, 2009). The *sub-1* (L*er*) was described previously (Chevalier *et al.*, 2005). The *sub-9* mutant (Col), carrying a T-DNA insertion (SAIL_1158_D09), was described in (Vaddepalli *et al.*, 2011). The *chc1-1* (SALK_112213), *chc1-2* (SALK_103252), *chc2-1* (SALK_028826) and *chc2-2* (SALK_042321) alleles (all Col) (Alonso *et al.*, 2003) were described in (Kitakura *et al.*, 2011). Wild-type and mutant plants were transformed with different constructs using Agrobacterium strain GV3101/pMP90 (Koncz and Schell, 1986) and the floral dip method (Clough and Bent, 1998). Transgenic T1 plants were selected on Kanamycin (50 µg/ml), Hygromycin (20 µg/ml) or Glufosinate (Basta) (10 µg/ml) plates and transferred to soil for further inspection. The hydroxytamoxifen-inducible line INTAM>>RFP-HUB/Col line (HUB) was described previously (Robert *et al.*, 2010; Kitakura *et al.*, 2011). Seedlings were grown on half-strength Murashige and Skook (1/2 MS) agar plates (Murashige and Skoog, 1962).

### Recombinant DNA work

For DNA and RNA work standard molecular biology techniques were used. PCR-fragments used for cloning were obtained using Q5 high-fidelity DNA polymerase (New England Biolabs, Frankfurt, Germany). All PCR-based constructs were sequenced. Primer sequences used in this work are listed in supplementary material Table SX.

### Reporter constructs

The pCAMBIA2300-based pSUB::SUB:EGFP construct was described previously (Vaddepalli *et al.*, 2011). To obtain pUBQ10::SUB:EGFP, a 2 kb promoter fragment of *UBQ10* (At4g05320) was amplified from L*er* genomic DNA using primers pUBQ(KpnI)_F and pUBQ(AscI)_R. The resulting PCR product was digested using KpnI/AscI and used to replace the pSUB fragment in pSUB::SUB:EGFP. The pGL2::GUS:EGFP construct was assembled using the GreenGate system (Lampropoulos *et al.*, 2013). The promoter region of *GL2* (AT1G79840) was amplified with primer pGL2 _F1 and pGL2_R1 from genomic Col-0 DNA. The internal BsaI site was removed during the procedure as described in (Lampropoulos *et al.*, 2013). The GUS coding sequence was amplified from plasmid pBI121 (Jefferson *et al.*, 1987) with primer GUS_F and GUS_R, digested with BsaI and used for further cloning.

### Genotyping PCR

PCR-based genotyping was performed with the following primer combinations: *sub-9*, SUB_LP158, SUB_RP158, and SAIL_LB2; *chc2 salk-042321*, CHC2-LP321, CHC2-RP321, and SALK_LBb1.3; *chc2 salk-028826*, CHC2_LP826, CHC2_RP826, and SALK_LBb1.3; *chc1 salk-112213*, CHC1_LP213, CHC1_RP213, and SALK_LBb1.3; *chc1 salk-103252*, CHC1_LP252, CHC1_RP252, and SALK_LBb1.3.

### Chemical treatments

Brefeldin A (BFA), cycloheximide, tyrphostin A23 (TyrA23), Wortmannin, Concanamycin A (ConcA) were obtained from Sigma-Aldrich and used from stock solutions in DMSO (50 mM BFA, cycloheximide, TyrA23; 30 mM Wortmannin, 2 mM ConcA). FM4-64 was purchased from Molecular Probes (2 mM stock solution in water). Five day-old seedlings were incubated for the indicated times in liquid 1/2 MS medium containing 50 µM BFA, 50 µM cycloheximide, 75 µM TyrA23, 33 µM Wortmannin, and 2 µM ConcA. For FM4-64 staining seedlings were incubated in 4 µM FM4-64 in liquid 1/2 MS medium for 5 minutes prior to imaging. 4-hydroxytamoxifen was obtained from Sigma-Aldrich (10 mM stock solution in ethanol). Seedlings were grown for 3 days on 1/2 MS plates, transferred onto 1/2 MS plates containing 2 µM 4-hydroxytamoxifen (or ethanol as mock treatment) for four days and then imaged using confocal microscopy.

### Immunoprecipitation and western blot analysis

500 mg of 7-day wild-type or transgenic seedlings were lysed using a TissueLyser II (Qiagen) and homogenized in 1 ml lysis buffer A (50mM Tris-HCl pH7.5, 100 mM NaCl, 0.1 mM PMSF, 0.5% Triton X-100, protease inhibitor mixture (Roche)). Cell lysate was mildly agitated for 15 min on ice and centrifuged for 15 minutes at 13000 g. For lines carrying GFP-tagged proteins supernatant was incubated with GFP-TRAP_MA magnetic agarose beads (ChromoTek) for 2 hours at 4°C. Beads were concentrated using a magnetic separation rack. Samples were washed four times in buffer B (50mM Tris-HCl pH7.5, 100 mM NaCl, 0.1 mM PMSF, 0.2% Triton X-100, protease inhibitor mixture (Roche)). Bound proteins were eluted from beads by heating the samples in 30 µl 2x Laemmli buffer for 5 minutes. Samples were separated by SDS-PAGE and analyzed by immunoblotting according to standard protocols. Primary antibodies included mouse monoclonal anti-GFP antibody 3E6 (Invitrogen/Thermo Fisher Scientific), mouse monoclonal anti-ubiquitin antibody P4D1 (Santa Cruz Biotechnology), and polyclonal anti-CHC antibody AS10 690-ALP (Agrisera). Secondary antibodies were obtained from Pierce/ThermoFisher Scientific: goat anti-rabbit IgG antibody (1858415) and goat anti-mouse IgG antibody (1858413).

### Microscopy and art work

To assess the cellular structure of floral meristems samples were stained with mPS-PI (Truernit *et al.*, 2008). Confocal laser scanning microscopy was performed with an Olympus FV1000 setup using an inverted IX81 stand and FluoView software (FV10-ASW version 01.04.00.09) (Olympus Europa GmbH, Hamburg, Germany) equipped with a water-corrected 40x objective (NA 0.9) at 3x digital zoom. For SUB:EGFP subcellular localization upon drug treatments or colocalization with endosomal markers confocal laser scanning microscopy was performed on epidermal cells of root meristems located about eight to 12 cells above the quiescent center using a Leica TCS SP8 X microscope equipped with GaAsP (HyD) detectors. The following objectives were used: a water-corrected 63x objective (NA 1.2), a 40x objective (NA 1.1), and a 20x immersion objective (NA 0.75). Scan speed was set at 400 Hz, line average at between 2 and 4, and the digital zoom at 4.5 (colocalization with FM4-64), 3 (drug treatments) or 1 (root hair patterning). EGFP fluorescence excitation was done at 488 nm using a multi-line argon laser (3 percent intensity) and detected at 502 to 536 nm. FM4-64 fluorescence was excited using a 561 nm laser (1 percent intensity) and detected at 610 to 672 nm. For the direct comparisons of fluorescence intensities, laser, pinhole, and gain settings of the confocal microscope were kept identical when capturing the images from the seedlings of different treatments. The intensities of fluorescence signals at the PM were quantified using Leica LAS X software (version 3.3.0.16799). For the measurement of the fluorescence levels at the PM optimal optical sections of root cells were used for measurements. On the captured images the fluorescent circumference of an individual cell (ROI, region of interest) was selected with the polygon tool. The mean pixel intensity readings for the selected ROIs were recorded and the average values were calculated. For determination of colocalization, the distance from the center of each EGFP spot to the center of the nearest FM4-64, mKO or mRFP signal was measured by hand on single optical sections using ImageJ/Fiji software (Schindelin *et al.*, 2012). If the distance between two puncta was below the resolution limit of the objectives lens (0.24 µm) the signals were considered to colocalize (Ito *et al.*, 2012). Arabidopsis seedlings were covered with a 22×22 mm glass cover slip of 0.17 mm thickness (No. 1.5H, Paul Marienfeld GmbH & Co. KG, Lauda-Königshofen, Germany). Scanning electron microscopy was performed essentially as reported previously (Schneitz *et al.*, 1997; Sieburth and Meyerowitz, 1997). Images were adjusted for color and contrast using ImageJ/Fiji software.

## Results

### The endocytic route of SUB:EGFP

To investigate the endocytic pathway followed by SUB we made use of a previously well-characterized line carrying the *sub-1* null allele and a transgene encoding a SUB:EGFP translational fusion driven by its endogenous promoter (pSUB::SUB:EGFP). The line exhibits a wild-type phenotype demonstrating the presence of a functional reporter (Vaddepalli *et al.*, 2011; Vaddepalli *et al.*, 2014). We studied the subcellular distribution of the pSUB::SUB:EGFP reporter signal in epidermal cells of the root meristem using confocal laser scanning microscopy. These cells serve as an ideal model as *SUB* promotes the early patterning of root hairs, cells that are generated by the epidermis (Dolan *et al.*, 1993). In the absence of any obvious exogeneous stimulation of SUB signaling we observed SUB:EGFP signal at the PM and in cytoplasmic foci (Fig. 1A). Moreover, we noticed that the SUB:EGFP signal labelled structures resembling vesicles as well as the vacuole. These observations raise the possibility that SUB:EGFP undergoes internalization from the PM and is shuttled to the vacuole for degradation.

**Figure 1.**
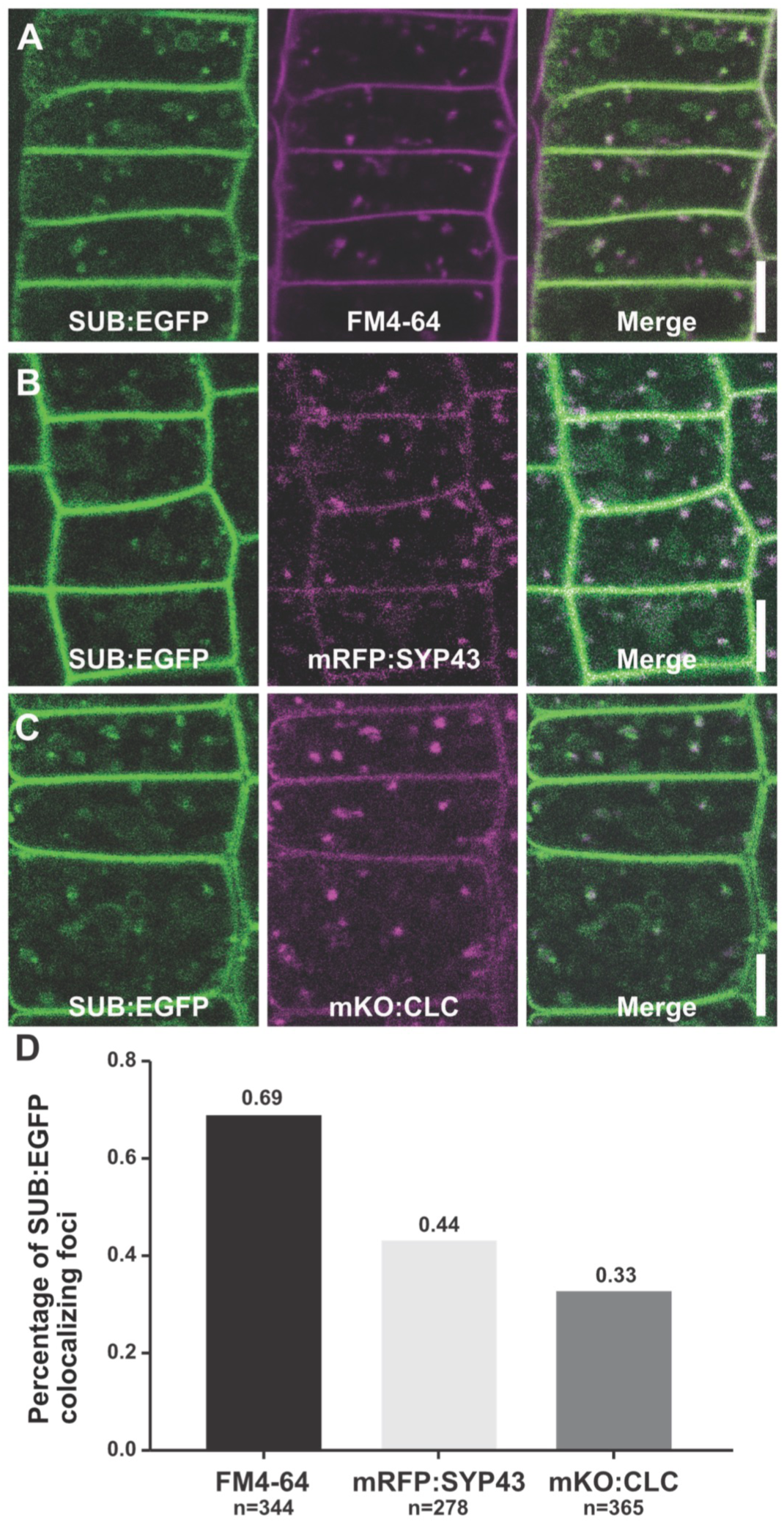
Sub-cellular localization of SUB:EGFP. Fluorescence micrographs show optical sections of epidermal cells of root meristems of five to six days-old seedlings. **(A)** Partial colocalization of SUB:EGFP and FM4-64 foci upon treating cells with FM4-64 for five minutes. **(B)** Partial colocalization of SUB:EGFP and mRFP:SYP43 puncta. **(C)** Partial colocalization of SUB:EGFP and mKO:CLC signals. **(D)** Quantitative colocalization analysis of SUB:EGFP-positive foci and reporter signals shown in **A**, **B** and **C**. *n*, total number of analyzed SUB:EGFP foci. Scale bars: 5 µm.

To assess the early process of SUB:EGFP endocytosis we imaged cells upon a 5-minutes treatment with the endocytic tracer dye FM4-64 (Fig. 1A, D). Using a previously described criterion for colocalization (Ito *et al.*, 2012) the internal SUB:EGFP and FM4-64 signals were considered colocalized when the distance between the centers of the two types of signals was below the limit of resolution of the objective, in our case 0.24 µm. We observed that 70 percent (n = 344) of all cytoplasmic SUB:EGFP foci were also marked by FM4-64 supporting endocytosis of SUB:EGFP. To further investigate internalization of SUB:EGFP we treated five days-old seedlings with Wortmannin. Wortmannin is a phosphatidylphosphate-3-kinase inhibitor that among others interferes with vesicle formation from the PM (Tse *et al.*, 2004; Wang *et al.*, 2009; Ito *et al.*, 2012; Cui *et al.*, 2016). We analyzed the number of internal SUB:EGFP-labelled puncta in cells upon treatment with 33 µM Wortmannin for 60 minutes. We found a substantial reduction in the number of such puncta in drug-treated cells when compared to mock-treated cells (Fig. 2A). Moreover, we noted a significant increase in SUB:EGFP signal intensity at the PM in Wortmannin-treated cells.

**Figure 2.**
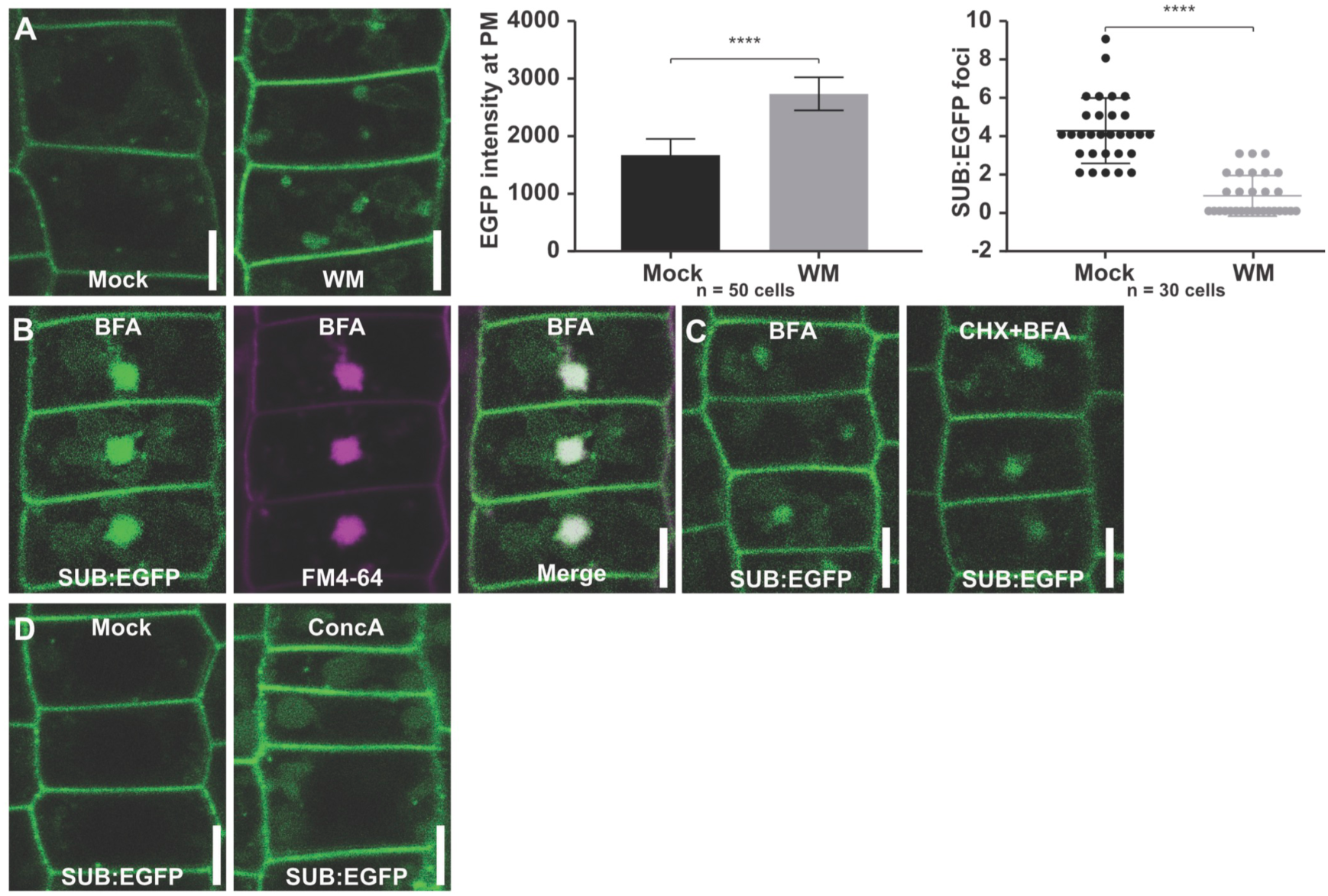
Sub-cellular localization of SUB:EGFP upon drug treatments. Fluorescence micrographs show optical sections of epidermal cells of root meristems of five to six days-old seedlings. **(A)** Subcellular localization of SUB:EGFP signal in the presence of Wortmannin and DMSO (mock control) (left). Graphs represent quantification of the EGFP intensity at plasma membrane (middle panel, n = 50 cells across six roots) and the number of SUB:EGFP-positive endosomes per cell (right panel, n = 30 cells across six roots) after incubation. Asterisks represent statistical significances (P<0.0001) as judged by Student’s *t* test. **(B)** SUB:EGFP signal is detected in BFA bodies upon BFA treatment. **(C)** SUB:EGFP signal is detected in BFA compartments in the presence of CHX. **(D)** SUB:EGFP signal is observed in lytic vacuoles after ConcA treatment. Abbreviations: ROI, region of interest. Scale bars: 5 µm.

To explore if endosomal trafficking of SUB:EGFP involves the TGN/EE we investigated colocalization of SUB:EGFP with the TGN marker mRFP:SYP43 (Ebine *et al.*, 2011; Ito *et al.*, 2012; Uemura *et al.*, 2012) (Fig. 1B, D). We observed a frequency of 44 percent colocalization (n = 278) between internal SUB:EGFP and RFP:SYP43 puncta. To further assess colocalization of SUB:EGFP with the TGN we made use of a previously characterized translational fusion between CLC2 and monomeric Kushiba Orange under the control of the cauliflower mosaic virus 35S promoter (mKO:CLC) (Fujimoto *et al.*, 2010). CLC2 fused to fluorescent tags also localizes to the TGN in live cell imaging experiments (Ito *et al.*, 2012). We observed a frequency of 33 percent colocalization (n = 365) between internal SUB:EGFP and mKO:CLC puncta (Fig. 1C, D). To corroborate the presence of SUB:EGFP at the TGN/EE we exposed *sub-1 pSUB::SUB:EGFP* seedlings to the fungal toxin Brefeldin A (BFA). Treatment with BFA results in the formation of so-called BFA compartments or bodies that contain secretory and endocytic vesicles (Robinson *et al.*, 2008; Paez Valencia *et al.*, 2016). We observed prominent SUB:EGFP signal in BFA compartments in root epidermal cells of seedlings treated with DMSO for 30 minutes followed by a DMSO/BFA (50 µM) treatment for 60 minutes, confirming previous data (Fig. 2B) (Kwak and Schiefelbein, 2008; Yadav *et al.*, 2008; Vaddepalli *et al.*, 2011; Wang *et al.*, 2016b).

To explore the relative contribution of signal at the TGN/EE originating from the secretion of newly translated SUB:EGFP versus endocytic SUB:EGFP-derived signal we first treated seedlings with the protein synthesis inhibitor cycloheximide (50 µM) for 60 minutes followed by co-incubation with 50 µM BFA for 30 minutes. In those seedlings SUB:EGFP still prominently localized to BFA bodies (Fig. 2C) as noted earlier (Wang *et al.*, 2016a). Taken together, the results indicate that a large fraction of SUB:EGFP in BFA bodies originated from the PM.

We next investigated if internalized SUB:EGFP is sorted into MVBs. Apart from affecting vesicle formation at the PM Wortmannin also interferes with the maturation of LEs and causes formation of enlarged MVB/LEs (Tse *et al.*, 2004; Wang *et al.*, 2009; Cui *et al.*, 2016). Treating seedlings for 60 minutes with 33 µM wortmannin results in the formation of large globular structures labelled by SUB:EGFP signal (Fig. 2A). Such structures are typical for enlarged MVBs (Jia *et al.*, 2013). In accordance with these results SUB:EGFP was detected at MVBs in immunogold electron microscopy experiments (Vaddepalli *et al.*, 2014).

Concanamycin A (ConcA) inhibits vacuolar ATPase activity at the TGN/EE and in the tonoplast thereby interfering with the trafficking of newly synthesized materials to the PM, the transport of cargo from the TGN/EE to the vacuole, and the vacuolar degradation of cargo (Dettmer *et al.*, 2006; Robinson *et al.*, 2008; Viotti *et al.*, 2010; Scheuring *et al.*, 2011). Upon treatment with 2 µM ConcA for 1 hour seedlings showed large roundish structures labelled by a diffuse SUB:EGFP signal (Fig. 2D) indicating that SUB:EGFP was not degraded efficiently and thus accumulated in the vacuole.

Taken together the results are consistent with the notion that the endocytic route of SUB:EGFP involves the TGN/EE, the MVB/LEs, and the vacuole where it becomes degraded. A noticeable portion of SUB:EGFP puncta colocalizes with the TGN/EE, supporting passage of SUB:EGFP through the TGN/EE. However, we cannot exclude that a fraction of SUB:EGFP also traffics via an TGN/EE-independent route, as does for example the AtPep1-PEPR1 signaling complex (Ortiz-Morea *et al.*, 2016).

### SUB:EGFP is ubiquitinated in vivo

Ubiquitination plays an important role in endocytosis and endosomal sorting of PM proteins (MacGurn *et al.*, 2012; Paez Valencia *et al.*, 2016; Isono and Kalinowska, 2017), such as the brassinosteroid receptor BRI1 (Martins *et al.*, 2015) or the auxin efflux facilitator PINFORMED 2 (PIN2) (Leitner *et al.*, 2012). To test if SUB:EGFP is ubiquitinated in vivo we made use of our *sub-1 pSUB::SUB:EGFP* reporter line as well as a previously described line carrying the SUB:EGFP translation fusion driven by the *UBIQUITIN10* (*UBQ*) promoter (pUBQ::SUB:EGFP) (Vaddepalli *et al.*, 2017). We immunoprecipitated SUB:EGFP from seven days-old, plate-grown seedlings using an anti-GFP antibody. Immunoprecipitates were subsequently probed with the commonly used P4D1 anti-ubiquitin antibody recognizing mono- and polyubiquitinated proteins. P4D1-dependent signal could not be reproducibly detected when testing immunoprecipitates from lines expressing the pSUB::SUB:EGFP reporter due to low abundance of SUB:EGFP in the immunoprecipitate. By contrast, we clearly observed a high-molecular weight smear in immunoprecipitates obtained from pUBQ::SUB:EGFP lines (Fig. 3). This smear is typical for ubiquitinated proteins. We did not detect signals in immunoprecipitates obtained from wild-type seedlings. The results indicate that a fraction of SUB proteins becomes ubiquitinated.

**Figure 3.**
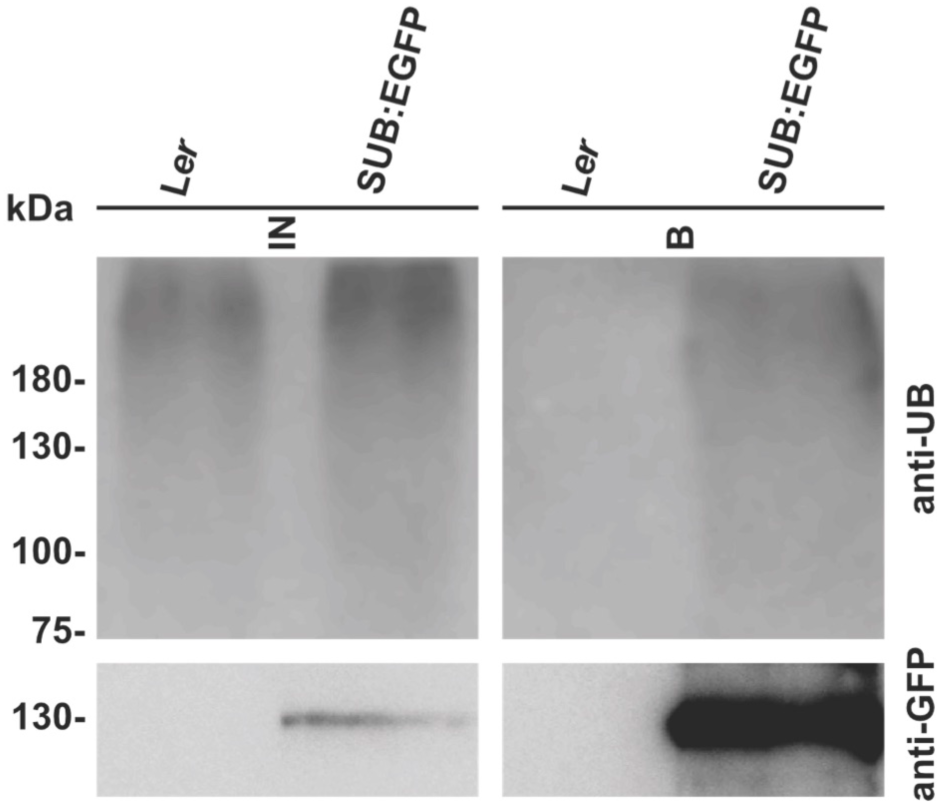
*In vivo* ubiquitination of SUB. Western blot analysis of immunoprecipitates obtained from wild type (L*er*) and *sub-1* pUBQ::gSUB:EGFP lines are shown. Immunoprecipitation was performed using an anti-GFP antibody. Immunoblots were probed with the P4D1 anti-Ub antibody (top panel) and an anti-GFP antibody (bottom panel). Abbreviations: B: bound fraction; IN, input.

### SUB:EGFP internalization involves clathrin-mediated endocytosis

So far, the obtained results indicate that SUB:EGFP is continuously internalized and eventually targeted to the vacuole for degradation. Next we wanted to assess if SUB:EGFP relates to a clathrin-dependent process. We first tested if SUB:EGFP and endogenous CHC occur in the same complex in vivo. To this end we immunoprecipitated SUB:EGFP from seven days-old, plate-grown *pUBQ::SUB:EGFP sub-1* seedlings using an anti-GFP antibody. Immunoprecipitates were subsequently probed using an anti-CHC antibody. We could detect a CHC-signal in immunoprecipitates derived from SUB:EGFP plants but not from wild-type (Fig. 4) indicating that SUB:EGFP and CHC are present in the same protein complex in vivo.

**Figure 4.**
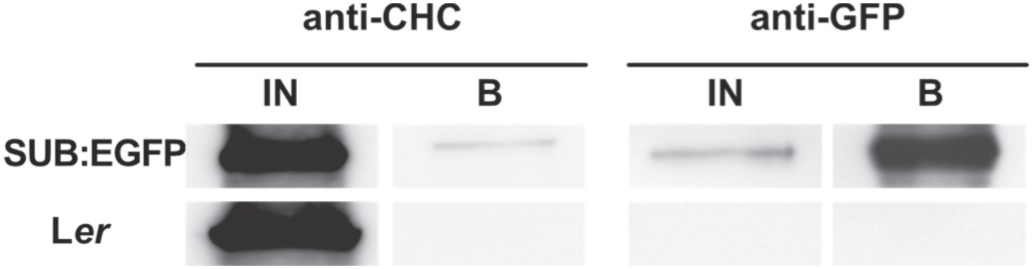
Co-immunoprecipitation of CHC with SUB:EGFP. Total extracts of seven day-old SUB:EGFP-expressing seedlings (upper panel) or wild-type seedlings (lower panel) were immunoprecipitated using GFP-Trap MA beads. Immunoblots were probed with anti-CHC (left panel) or anti-GFP antibodies (right panel). Abbreviations: B: bound fraction; IN, input.

Next we assessed the contribution of clathrin to the internalization and subcellular distribution of SUB:EGFP. To this end, we investigated the effects of a transient but robust impairment of clathrin activity on the internalization and subcellular distribution of SUB:EGFP. Ectopic expression of the C-terminal part of CHC1 (HUB1) results in a dominant-negative effect due to the HUB1 fragment binding to and out-titrating clathrin light chains (Liu *et al.*, 1995). To assess the effect of the presence of the HUB fragment on the subcellular distribution of SUB:EGFP we crossed a previously characterized 4-hydroxytamoxifen-inducible INTAM>>RFPCHC1 (HUB) line (Robert *et al.*, 2010; Kitakura *et al.*, 2011) into a Col-0 wild-type line carrying the pUBQ::SUB:EGFP reporter. We then analyzed epidermal cells of the root meristem of HUB/pUBQ::SUB:EGFP plants, hemizygous for each transgene, upon induction.

We first determined the length of induction period that enabled us to detect by confocal microscopy a defect in endocytosis, as indicated by a reduction of internal FM4-64 foci following a 5 to 10 minutes exposure to the stain. Under our growth conditions a significant reduction of internal FM4-64 puncta was observed after three days of continuous growth on induction medium while near complete absence of internal FM4-64 foci was detected after four days (Fig. 5B, C). If SUB:EGFP participates in CME a block in HUB-sensitive endocytosis should result in fewer internal SUB:EGFP-labelled foci and higher SUB:EGFP signal at the PM when compared to the SUB:EGFP-derived signal of a control line. We found a significant reduction in cytoplasmic SUB:EGFP puncta in the HUB/pUBQ::SUB:EGFP line after three days of growth on induction medium in comparison to the control (Fig. 5B). Upon four days of induction we detected an increase in SUB:EGFP signal at the PM (Fig. 5C). Taken together, our results suggest that CME contributes to the internalization of SUB:EGFP.

**Figure 5.**
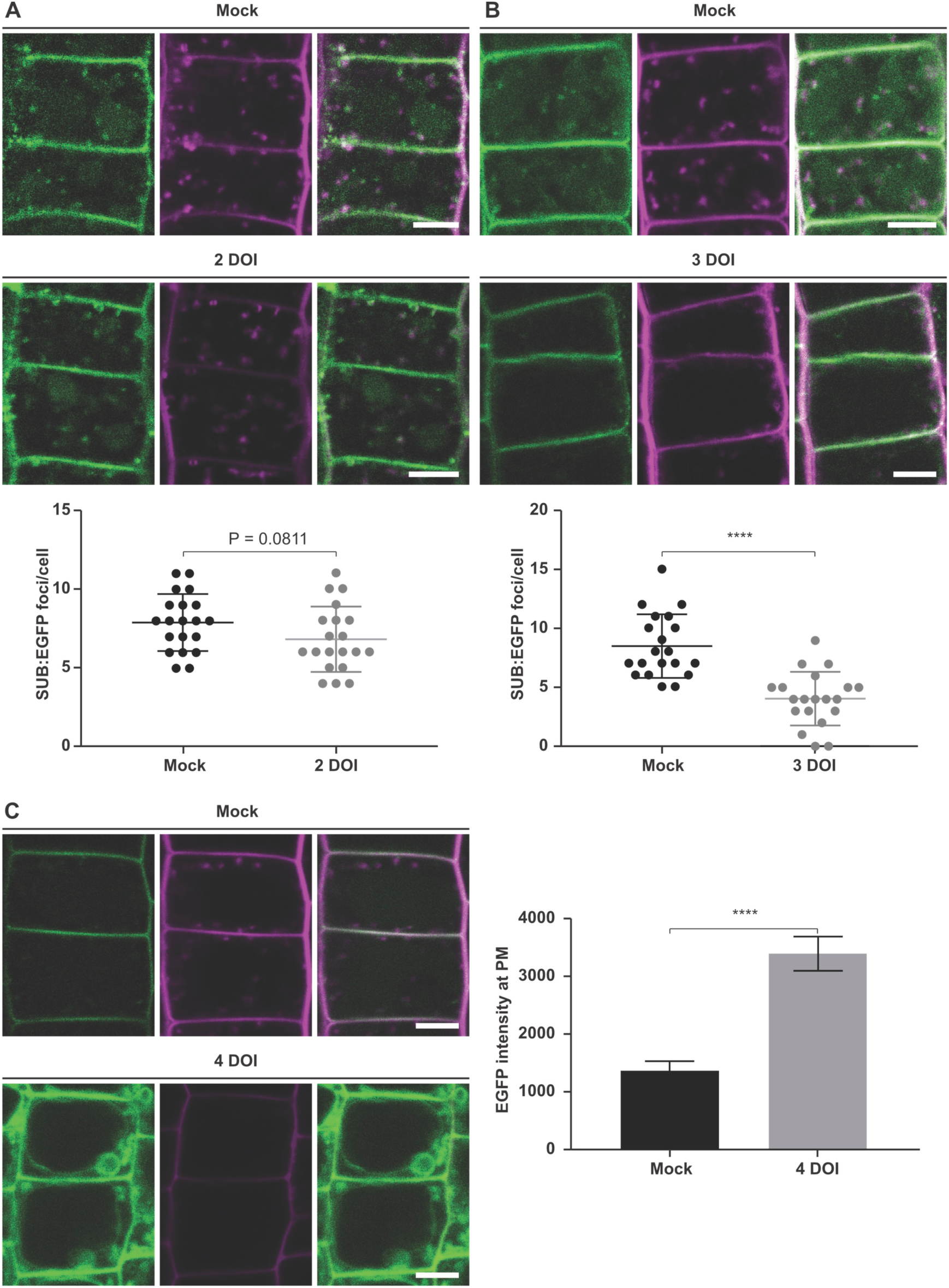
Requirement of clathrin function for SUB endocytosis. Fluorescence micrographs show optical sections of epidermal cells of root meristems. **(A) to (C)** Internalization of SUB:EGFP and uptake of endocytic tracer dye FM4-64 in epidermal meristems cells of three days-old INTAM>>RFP-CHC1 (HUB1)/pUBQ::SUB:EGFP seedlings that were placed on 2 µM 4-hydroxytamoxifen-containing induction medium for two, three, or four days, respectively. Ethanol served as mock. Graphs represent quantification of the number of SUB:EGFP-positive spots per cell **(A, B)** and of the EGFP intensity at plasma membrane **(C)** after incubation. Abbreviation: DOI, days on induction medium. Scale bars: 5 µm.

### SUB genetically interacts with CLATHRIN HEAVY CHAIN

To further assess the role of clathrin in the SUB signaling mechanism we tested a possible genetic interaction between *SUB* and *CHC*. To this end we made use of several previously characterized T-DNA insertion lines carrying knock-out alleles of *CHC1* and *CHC2* (Kitakura *et al.*, 2011). Plants lacking *CHC1* as well as *CHC2* function appear to be lethal (Kitakura *et al.*, 2011). However, mutations in individual *CHC* genes result in endocytosis defects and affect for example polar distribution of PIN proteins, internalization of ATRBOHD, stomatal movement, and resistance to powdery mildew (Kitakura *et al.*, 2011; Hao *et al.*, 2014; Wu *et al.*, 2015; Larson *et al.*, 2017).

To test if clathrin is involved in *SUB*-controlled processes we first investigated if *chc1* and *chc2* mutants show a defect in root hair patterning. To this end we generated homozygous *chc* mutants carrying a translational fusion of bacterial ß-glucuronidase (GUS) to EGFP (GUS:EGFP) under the control of the Arabidopsis *GLABRA2* (*GL2*) promoter (pGL2::GUS:EGFP). The *GL2* promoter drives expression specifically in non-root hair cells and is commonly used to monitor root hair patterning (Masucci *et al.*, 1996; Kwak *et al.*, 2005). Interestingly, we found that all *chc* alleles tested showed root hair patterning defects similar to *sub-9* with *chc2* alleles causing more prominent aberrations compared to *chc1* mutations (Fig. 6, Supplementary Fig. S1). In addition, *chc1 sub-9* or *chc2 sub-9* double mutants did not show an obviously exacerbated phenotype indicating that *CHC1*, *CHC2* and *SUB* do not act in an additive fashion. Thus, the results indicate that *CHC1* and *CHC2* promote root hair pattern formation and that they function in the same genetic pathway than *SUB*.

**Figure 6.**
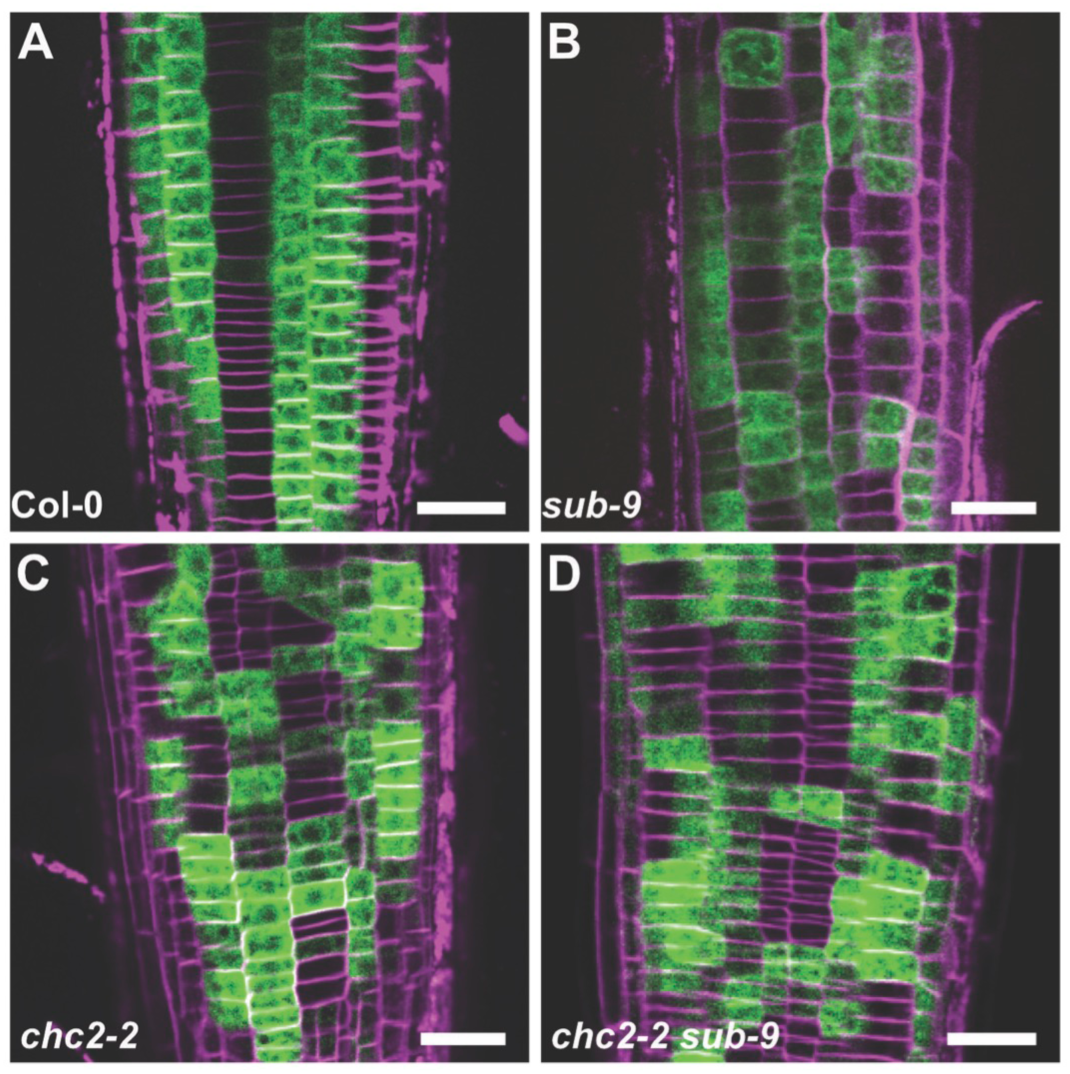
Expression pattern of the pGL2::GUS:EGFP reporter in *chc2-2* and *chc2-2 sub-9* mutants. Fluorescence micrographs show optical sections of epidermal cells of root meristems of seven days-old seedlings. FM4-64 was used to label cell outlines. **(A)** Col-0. **(B)** *sub-9*. **(C)** *chc2-2*. **(D)** *chc2-2 sub-9*. Note the similarly altered pattern in **(B)** to **(D)**. Scale bars: 25 µm.

Next, we assessed if *CHC1* and *CHC2* participate in *SUB*-dependent floral development. In the Col-0 background null alleles of *SUB* cause a weaker floral phenotype when compared to similar alleles in the L*er* background (Vaddepalli *et al.*, 2011). Still, the *sub-9* allele causes mild silique twisting, mis-oriented cell division plants in the L2 layer of floral meristems, and ovule defects (Fig. 7) (Tables 1 and 2) (Vaddepalli *et al.*, 2011). By contrast, we did not detect any obvious defects in floral meristems, flowers and ovules of plants homozygous for the tested *chc1* or *chc2* alleles (Fig. 7, Supplementary Fig. S2) (Tables 1 and 2). We then investigated the phenotype of *chc1 sub-9* and *chc2 sub-9* double mutants. Interestingly, the cell division defects in the L2 layer of the FM were reduced in *chc1 sub-9* and *chc2 sub-9* double mutants in comparison to *sub-9* single mutants (Fig. 7, Supplementary Fig. S2) (Table 1). Suppression of the *sub-9* phenotype in *chc1 sub-9* or *chc2-sub-9* double mutants was also observed for silique twisting and ovule development (Fig. 7, Supplementary Fig. S2) (Table 2). The results suggest that *SUB* is a negative genetic regulator of *CHC1* and *CHC2* function in floral meristem, ovule and silique development.

**Figure 7.**
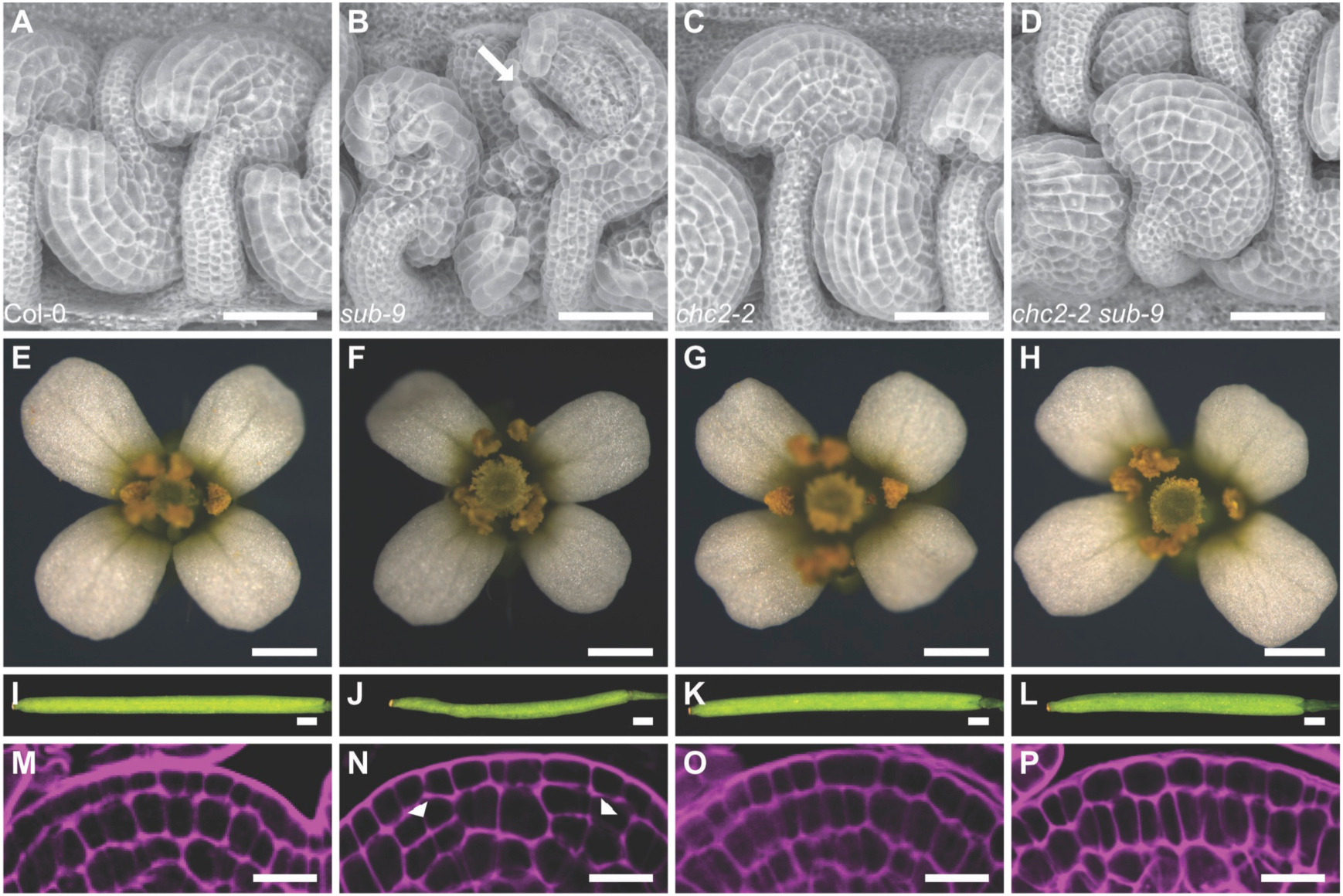
Phenotype comparison between Col-0, *sub-9*, *chc2-2* and *chc2-2 sub-9*. **(A)** to **(D)** Scanning electron micrographs of stage 4 ovules (stages according to (Schneitz *et al.*, 1995)). **(B)** Note the aberrant outer integument (arrow). **(E)** to **(H)** Morphology of mature stage 13 or 14 flowers (stages according to (Smyth *et al.*, 1990)). **(I)** to **(L)** Morphology of siliques. **(M)** to **(P)** Central region of stage 3 floral meristems stained with pseudo-Schiff propidium iodide (mPS-PI). **(N)** Arrowheads indicate aberrant cell division planes. **(P)** Note the defects of the *sub-9* phenotype were partially rescued in *chc sub-9* double mutants. Scale bars: **(A)** to **(D)** 50 µm, **(E)** to **(H)** 0.5 mm, **(I)** to **(L)** 1 mm, **(M)** to **(P)** 10 µm.

**Table 1.** Number of periclinal cell divisions in the L2 layer of stage 3 floral meristems.

| Genotype | NPCD<sup>a</sup> | Percentage | NFM<sup>b</sup> |
| --- | --- | --- | --- |
| Col | 12 | 17.6 | 68 |
| *sub-9* | 17 | 36.2 | 47 |
| *chc1-1* | 6 | 23.1 | 26 |
| *chc1-2* | 7 | 20.0 | 35 |
| *chc2-1* | 7 | 22 | 31 |
| *chc2-2* | 5 | 22.5 | 25 |
| *chc1-1 sub-9* | 6 | 20.0 | 30 |
| *chc1-2 sub-9* | 7 | 19.4 | 36 |
| *chc2-1 sub-9* | 7 | 25.9 | 27 |
| *chc2-2 sub-9* | 7 | 18.9 | 37 |

<sup>a</sup>Number of periclinal cell divisions observed
<sup>b</sup>Number of floral meristems observed

**Table 2.** Comparison of integument defects between *sub-9*, *chc* and *chc sub-9* mutants.

| Genotype | N total | N with defects | Percentage |
| --- | --- | --- | --- |
| Col | 274 | 0 | 0 |
| *sub-9* | 291 | 82 | 28.2 |
| *chc1-1* | 130 | 0 | 0 |
| *chc1-2* | 121 | 0 | 0 |
| *chc2-1* | 126 | 0 | 0 |
| *chc2-2* | 230 | 0 | 0 |
| *chc1-1 sub-9* | 235 | 14 | 6 |
| *chc1-2 sub-9* | 185 | 11 | 6 |
| *chc2-1 sub-9* | 211 | 14 | 6.6 |
| *chc2-2 sub-9* | 237 | 13 | 5.5 |

## Discussion

An impressive body of published work has elucidated many of the intricacies of receptor-mediated endocytosis of plant RKs. Much is known about the internalization and endocytic trafficking of plant RKs with functional kinase domains. The atypical RK SUB carries an inconspicuous kinase domain, however, enzymatic kinase activity could not be demonstrated in in vitro biochemical experiments and is not required for its function in vivo (Chevalier *et al.*, 2005; Vaddepalli *et al.*, 2011; Kwak *et al.*, 2014). Using SUB as a model we have explored the endocytic route of an atypical RK. We investigated this process by examining the subcellular distribution of a functional SUB:EGFP reporter in epidermal cells of the root meristem. Our data are compatible with the notion that PM-localized SUB becomes internalized and traffics from the TGN/EE to MVB/LEs and eventually the vacuole where it is destined for degradation. SUB:EGFP was observed to enter this endocytic route in the apparent absence of activation of SUB signaling by artificial stimulation or application of exogenous ligand. A similar observation was for example made for ACR4 (Gifford *et al.*, 2005). One interpretation of this finding could be that endogenous SUB ligand is always present in sufficient levels to promote SUB endocytosis. In another possible scenario, the rate of SUB internalization may be independent from ligand availability as was shown for BRI1 (Russinova *et al.*, 2004; Geldner *et al.*, 2007). In any case, our data indicate that the endocytic route of the atypical RK SUB for the most part seems to adhere to the established pattern of plant receptor-mediated endocytosis.

Apart from being a central signal for proteasome-mediated degradation ubiquitination is a major endocytosis determinant of PM proteins (Haglund and Dikic, 2012; Isono and Kalinowska, 2017). Several plant RKs are known to be ubiquitinated, including FLS2 (Lu *et al.*, 2011), BRI1 (Martins *et al.*, 2015; Zhou *et al.*, 2018), and LYK5 (Liao *et al.*, 2017). The observed in vivo ubiquitination of SUB:EGFP is compatible with the notion of SUB being internalized and transported to the vacuole for degradation. However, it remains to be determined which E3 ubiquitin ligase promotes ubiquitination of SUB and how SUB endocytosis relates to the control of its signaling. Internalization can be linked with downstream responses, as was demonstrated for FLS2 or the *At*Pep1-PEPR complex (Mbengue *et al.*, 2016; Ortiz-Morea *et al.*, 2016), or contribute to signal downregulation, as it is the case for BRI1 (Irani *et al.*, 2012; Zhou *et al.*, 2018) or LYK5 (Liao *et al.*, 2017).

Several lines of evidence support the notion of SUB:EGFP undergoing CME. First, CHC in vivo co-immunoprecipitated with SUB:EGFP. Second, we observed a reduction in intra-cellular SUB:EGFP puncta accompanied with a stronger SUB:EGFP signal at the PM in the HUB-line upon induction. Third, our genetic analysis revealed a connection of SUB with a clathrin-dependent process. Plants with a defect in *CHC2* show significantly reduced endocytosis rate of FM4-64 and aberrant polar localization of the polar auxin transporter PINFORMED 1 (PIN1) (Kitakura *et al.*, 2011) as well as reduced internalization of, for example, PEP1 {Ortiz-Morea et al., 2016, #6940}, FLS2 (Mbengue *et al.*, 2016), and BRI1 (Wang *et al.*, 2015). Accordingly, *chc2* mutants show multiple defects, including patterning defects in the embryo (Kitakura *et al.*, 2011), impaired mitogen-activated protein kinase (MAPK) activation (Ortiz-Morea *et al.*, 2016), and defective stomatal closure and callose deposition upon bacterial infection (Mbengue *et al.*, 2016). Our genetic analysis revealed that *CHC2,* and to a lesser effect *CHC1,* also affects root hair patterning. Importantly, it provides evidence for a biologically relevant interaction between *SUB* and a *CHC*-dependent process.

Interestingly, the data suggest that the type of genetic interaction between *SUB* and *CHC* depends on the tissue context. In the root, *SUB* and *CHC* promote root hair patterning. Several hypotheses are conceivable that could explain the result. As our data support the notion of SUB:EGFP undergoing CME, one model states that CME of SUB is required for root hair patterning. Therefore, SUB internalization in single *chc* mutants would be reduced resulting in a hyperaccumulation of SUB at the PM. Two alternative further scenarios are compatible with this notion. In the first scenario hyperaccumulation of SUB at the PM interferes with root hair patterning. This view is supported by the observation that not just a reduction of *SUB* activity but also ectopic expression of *SUB* in *p35S::SUB* plants results in a weak defect in root hair patterning (Kwak and Schiefelbein, 2007), similar to what we observed for *chc2* mutants. In the second scenario, a reduction of SUB internalization leads to fewer SUB-labelled endosomes, which in turn impairs root hair patterning. This scenario implies that SUB signals while being present on endosomes. In another model, a reduction of CHC activity could influence clathrin-dependent secretion of newly translated and/or recycled SUB to the PM thereby reducing the level of active SUB at the PM below a certain threshold. Finally, given the pleiotropic phenotype of *chc* mutants the genetic data do not rule out a more indirect interaction between *SUB* and *CHC*. Further work remains to be done to discriminate between the different possibilities. However, we currently favor the notion that CME of SUB is critical for root hair patterning as a block of CME of SUB:EGFP in the HUB line results in a reduction of internalized SUB:EGFP vesicles and elevated levels of SUB:EGFP at the PM.

In contrast to the positive genetic role of *SUB* and *CHC* in root hair patterning the apparent wild-type appearance of floral organs of *sub chc* double mutants indicates that *SUB* is a negative regulator of a *CHC*-dependent process during floral development. The molecular mechanism remains to be investigated. CME could for instance promote the internalization of a PM-resident signaling molecule, thereby attenuating its activity. This endocytic process could be counteracted upon by SUB. For example, in a *sub* mutant the activity of the hypothetical signaling factor at the PM is reduced through increased endocytosis. In a *sub chc* double mutant the principally higher level of internalization caused by the lack of SUB activity is offset by reduced CME due to impaired CHC function. Thus, the individual *sub* and *chc* effects cancel each other out in *sub chc* double mutants and the respective plants show flowers with apparent wild-type morphology. It will be an exciting challenge to unravel the molecular details of how SUB and clathrin interact to allow tissue morphogenesis in future studies.

## Supplementary data

Supplementary Figure S1: Comparison of the root hair patterning phenotype of wild type, *sub-9*, *chc1-1* and *chc2-1* mutants.

Supplementary Figure S2: Comparison of the floral phenotype of wild type, *sub-9*, *chc1-1* and *chc2-1* mutants.

Supplementary Table S1: List of all primers used in this study.

## Acknowledgements

We thank Jiří Friml for the INTAM>>RFP-CHC1 (HUB) line, Masaru Fujimoto for the mKO:CLC2 reporter and Tomohiro Uemura for the mRFP:SYP43 line. We also thank Silke Robatzek for critical reading of the manuscript. This work was funded by the German Research Council (DFG) through grant SFB924 (TP A06) to EI and SFB924 (TP A02) to KS.

